# Divergent selection for feed efficiency in pigs altered the duodenum transcriptome and DNA methylation profiles, resulting in greater responses to feed intake in the most efficient line

**DOI:** 10.1101/2022.11.03.515018

**Authors:** Guillaume Devailly, Katia Fève, Safia Saci, Julien Sarry, Sophie Valière, Jérôme Lluch, Olivier Bouchez, Laure Ravon, Yvon Billon, Hélène Gilbert, Juliette Riquet, Martin Beaumont, Julie Demars

## Abstract

Feed efficiency is a trait of interest in pigs as it contributes to lowering the ecological and economical costs of pig production. A divergent genetic selection experiment from a Large White pig population was performed for 10 generations, leading to pig lines with relative low-(LRFI, more efficient) and high-(HRFI, less efficient) residual feed intake (RFI). Feeding behaviour and metabolic differences have been previously reported between the two lines. We hypothesised that part of these differences could be related to differential sensing and absorption of nutrients in the proximal intestine.

Here we investigated the duodenum transcriptome and DNA methylation profiles comparing overnight fasting with ad libitum feeding in LRFI and HRFI pigs (n=24). We identified 1,106 differentially expressed genes between the two lines, notably affecting pathways of the transmembrane transport activity and related to mitosis or chromosome separation. The LRFI line showed a greater transcriptomic response to feed intake, with 2,222 differentially expressed genes before and after a meal, as compared to 61 differentially expressed genes in the HRFI line. Feed intake affected genes from both anabolic and catabolic pathways in the pig duodenum, such as rRNA production and autophagy. We noted that several nutrient transporter and tight junction genes were differentially expressed between lines and/or by short term feed intake. We also identified 409 differentially methylated regions in the duodenum mucosa between the two lines, while this epigenetic mark was less affected by feeding. However most of the differentially expressed genes were not associated with proximal changes in DNA methylation profiles.

Altogether, our findings highlighted that the genetic selection for feed efficiency in pigs changed the transcriptome profiles of the duodenum, and notably its response to feed intake, suggesting key roles for this proximal gut segment in mechanisms underlying feed efficiency.

## Introduction

In monogastric livestock, feed efficiency is the ability to convert the greater part of ingested feed into body weight. It is a complex trait with many known influencing factors such as nutrition, metabolism, genetics, microbiome, meteorological conditions, sanitary status, and gut microbiota. Feed efficiency is a trait of great interest for livestock since farming feed cost is a large part of the production costs. Feed also constitutes a large part of the environmental impacts of monogastric farming^1^. In addition to research into non-human-edible feed, improvement in the animal feed efficiency will reduce the amount of feed needed to raise livestock, thus contributing to the reduction of its environmental footprint^2–4^.

Feed efficiency can be measured as Residual Feed Intake (RFI): the difference between the amount of feed one animal is consuming and the amount predicted for its maintenance and production requirements via multiple regression on several traits (average metabolic body weight, average daily gain, and indicators of body composition). A negative RFI means that the animal is eating less (relatively high efficiency), and a positive RFI means that the animal is eating more (relatively low efficiency) than the population average for their specific growth rate and body composition. RFI is a heritable trait in pigs^5^. Differences in blood metabolome and brain, duodenum, liver, and muscle transcriptomes have previously been observed in pigs with contrasted feed efficiency^6–10^.

Gilbert et al. created a genetic divergent selection experiment from a Large White pig breed nucleus, resulting in two pig lines of relative low (LRFI) and high (HRFI) feed efficiency^5^. The response to selection has led to multiple genetic and genomic changes between the lines^11^. A previous transcriptomic study has compared muscle, adipose tissues and liver transcriptome from generation 8 of selection^12^, with pathways involved in immune response, response to oxidative stress and protein metabolism differentiating the two lines. These two lines also differ on their blood and muscle metabolism^13,14^—with a lower insulin level in the blood of LRFI pigs—and in their faecal microbiota^15^. The selection has led to distinct feeding behaviour between the lines: pigs from the LRFI line have a lower daily feed intake, eat more per visit to the feeder, stay longer in the feeder, eat faster, and wait longer between two visits than pigs from the HRFI line^16^.

The intestine is known to quickly adapt to various feeding challenges^17^. The duodenum is one of the key organs involved in nutrient sensing and satiety regulation, in relation with its proximal position in the gut^18,19^. We hypothesised that divergent selection for feed efficiency might have altered the duodenum physiological response to feed intake. The effect of feeding challenges on intestinal transcriptomes has been investigated by several groups. Zhang et al. reported that pigs submitted to different feeding frequencies, 1, 3 or 5 meals per day, have a different ileal and colonic mucosal transcriptome^20^, although they only investigated transcriptomes from overnight fasted pigs. Mazurais et al. studied the impact of a 2-day fasting in the jejunum expression levels of targeted genes in piglets, and noted 146 affected over the 954 studied genes^21^. In mice, Yoshioka et al. have investigated the effect of a high-fat and low-fat diet on the duodenal mucosa compared to fasted mice^22^. They noted only a modest effect on the transcriptome, but highlighted the downregulation of *Slc5a1* after feed intake. *Slc5a1* is the gene encoding for the sodium/glucose cotransporter 1 (SGLT1). It is also documented that the gene *Slc15a1* encoding for the peptide transporter 1 (PEPT1) is downregulated by feed intake in mice^23^. More generally, the Solute Carrier (SLC) gene family encodes transmembrane nutrient transporter, and is of great interest to better understand the digestive physiology^24–26^.

DNA methylation is an epigenetic mark playing important roles in cellular differentiation and in the regulation of gene expression^27,28^, including in the intestine^29^. DNA methylation is essential for proper intestinal epithelial cell differentiation^30^, and DNA methylation profiles change through cellular^31^ and developmental^32^ maturation. In cattle, genomic predictions for feed efficiency are correlated with DNA methylation differences at some imprinted loci^33^. A 36h fasting also resulted in changes in DNA methylation in adipose tissues in human^34^.

We hypothesised that differences between LRFI and HRFI pigs in the intestinal responses to feed intake might partially explain several of the physiological differences previously reported. We therefore investigated the transcriptomic and DNA methylation profiles of the duodenum mucosa before or after a meal in pigs from the 10th generation of LRFI and HRFI lines, in a total of 24 animals.

## Material & methods

### Animal production and sampling

This experimentation was authorised by the French Minister of Higher Education, Research and Innovation under the number APAFIS#21107-2018120415595562 v10, after examinations by the animal experimentation ethic committee number 084.

Animals were raised using standard care in the INRAE pig experimental facility GENESI^35^ up until the day before the procedure. Animals were from two French Large White lines of pigs that have been divergently selected on residual feed intakes for 10 generations^5^. Pigs were from three litters in each line, and of balanced sexes within litter when possible (one litter was represented by 3 males and one female). At weaning (28 days old on average), animals were split into 2 pens of 12 animals, full-sibs and sexes being equally distributed in the two pens (figure 1A). Pigs were slaughtered on the same day, at the age of 61 days old (min 60 days, max 62 days). The day before sampling, feed access was removed at 5 p.m. in both pens. Animals had free access to water. At 8 a.m. the next day, feed access was introduced back in one pen, but not in the other. Animals were slaughtered by electro-narcosis between 8.50 a.m. and 11.46 a.m., starting with animals left without feed access until 10.10 am where animals with unlimited feed access were also sampled (figure 1B, supplementary table 1). The gastro-intestinal tract was removed, and a 5 cm long section of the duodenum was sampled, opened longitudinally, and the mucosa was collected by scratching the internal duodenal section with a glass microscope slide. Samples were rinsed in PBS, and flash frozen in liquid nitrogen. Samples were preserved at -80°C until extraction.

**Figure 1:**
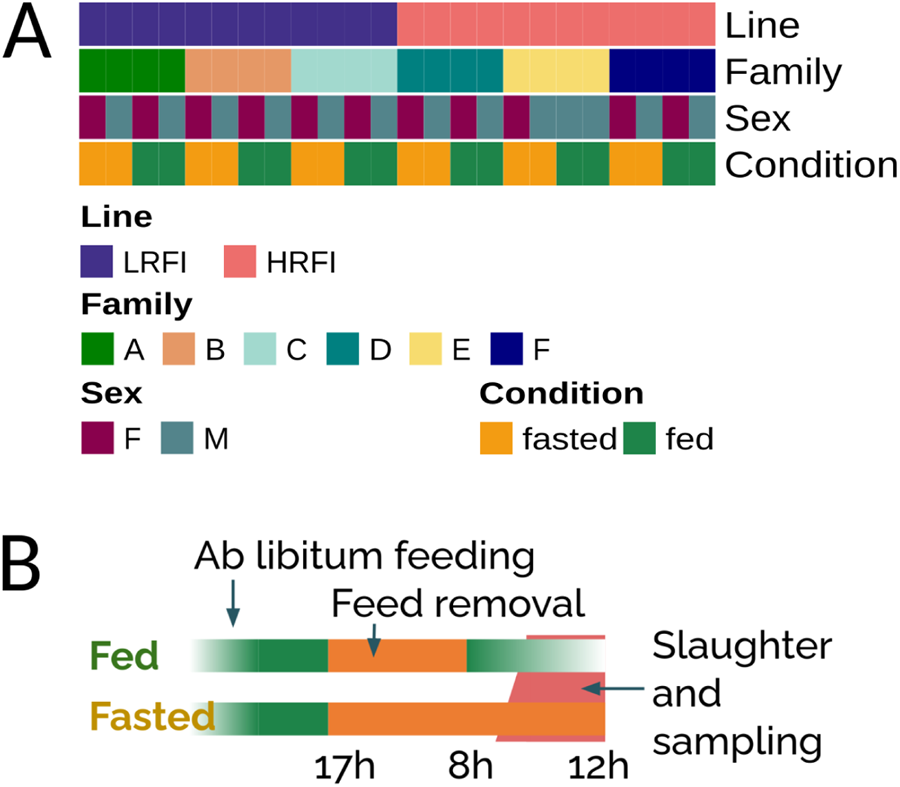
Experimental setup. **A**. 24 pigs from two pig lines (12 LRFI and 12 HRFI) were used, distributed in 3 litters of 4 pigs for each line, with two males and two females in each litter (to the exception of litter E). Half of the pigs were fasted while the other half was fed before sampling. **B**. Fasting procedure. Pigs were slotted into two pens with ad libitum feeding. Feed was removed from the two pens overnight, and reintroduced in the morning into one of the pens 2 hours before sampling started in this pen.

### Sample extraction, library preparation, and sequencing

Frozen duodenum samples were reduced to a fine powder with the mixer-mill MM400 by rapid agitation for 1 minute at 30Hz in a liquid-nitrogen cooled container with stainless steel beads. Fine powder samples were conserved at -80°C until extraction. DNA and RNA were extracted from the powder using NucleoSpin TriPrep mini kit columns (Macherey-Nagel), according to the manufacturer’s main instructions but with a few modifications. About 30-50 mg of powder were resuspended in 750 μL of RP1 buffer with 7.5μL of ß-mercaptoethanol then homogenised with the mixer-mill MM400 by rapid agitation for 2x2 minutes at 30Hz with stainless steel beads and incubated 10 minutes at room temperatures. Lysates were loaded into columns in two steps. DNA elution was performed using 65 μL of DNA elute solution. On-column digestion of residual DNA was performed for 30 minutes at room temperature. RNA was eluted using 100 μL of RNase-free water.

RNA-seq libraries were prepared according to Illumina’s protocols using the Illumina TruSeq Stranded mRNA sample prep kit to analyse mRNA. Briefly, mRNA were selected using poly-T beads. RNA were reverse transcribed to generate double stranded cDNA, then fragmented, and adaptors were ligated for sequencing. Eleven cycles of PCR were applied to amplify libraries. Library quality was assessed using a Fragment Analyser and libraries were quantified by qPCR using the Kapa Library Quantification Kit. RNA sequencing was performed on an Illumina HiSeq3000 using a paired-end read length of 2 x 150 pb with the Illumina HiSeq3000 Reagent Kits, at the GeT-PlaGe core facility^36^, INRAE Toulouse.

Four DNA samples (one female and one male LRFI, one female and one male HRFI) were artificially methylated using the M.Sss1 Enzyme (New England BioLabs) following the manufacturer instruction. DNA libraries (4 fully methylated control, 4 input controls, and 24 samples) were prepared following Bioo scientific’s protocol using the Bioo scientific NEXTflex Methyl-seq Library Prep Kit for Illumina Sequencing (part number NOVA-5118-02). Briefly, DNA was fragmented by sonication on a covaris M220 (400-500 bp), size selection was performed using AMPure XP beads and adaptors were ligated to be sequenced. Library preparation failed for one sample (an LRFI fed sample). The 4 fully methylated libraries and the 23 other samples were mixed at equimolarity when possible (62 ng of DNA per sample, otherwise taking a maximal volume of 20 μL resulting in 5-21 ng of DNA for four samples), and a single methyl-DNA precipitation (MeDP) was performed on 1 μg of the pool using Invitrogen’s Methylminer kit following the manufacturer instructions. After additions of the 4 inputs to the pool, 10 cycles of random PCR were performed on the mixed library. Library quality was assessed using an Agilent Fragment Analyser and the pool was quantified by qPCR using the Kapa Library Quantification Kit. MeDP sequencing (MeDP-seq) was performed on two lanes of an Illumina NovaSeq 6000 using a paired-end read length of 2 x 100 bp with the Illumina NovaSeq Reagent Kits S2, at the GeT-PlaGe core facility^36^, INRAE Toulouse.

RNA-seq and MeDP-seq libraries were demultiplexed, and the resulting *fastq* files are available at the ENA database under the id PRJEB46060 / ERP130249.

### RNA-seq bioinformatic processing

RNA-seq reads were processed using the nf-core/rnaseq pipeline^37^ (version 3.0), using Salmon^38^ pseudo-alignment quantifications, Ensembl reference genome Sscrofa11.1, and the corresponding gene annotation file from version 102 of Ensembl^39^. Normalised counts were processed using {tximports}^40^ to generate transcript per million (TPM) values (supplementary table 2), and gene-length normalised counts (supplementary table 3) were used to carry over differential gene expression analysis with limma voom^41^, using contrast matrices to test for different factors, and a false discovery rate (FDR) threshold of 5%. The pig line (LRFI or HRFI), condition (fed or fasted) and sex (castrated male or female) were used to build a model matrix. Contrast matrices were constructed to compare the two lines, and to compare the feeding effect within each line. A total of 13,738 genes were analysed, others having less than a total of 8 counts across our dataset. Gene ontology enrichment analysis was performed using {clusterProfiler}^42^, using Sus scrofa Gene Ontology. It should be noted that not all genes annotated in the pig genome are associated with a Gene Ontology term, leading to slightly lower gene numbers in gene lists used for the Gene Ontology Analyses. The set of genes used for differential gene expression analysis was used as a reference set. Enriched Gene Ontology were simplified for readability using the `simplify()` function, and simplified enriched ontologies were clustered using the `pairwise_termsim()` function. Heatmaps were generated using {ComplesHeatmap}^43^, and upset plots were generated using the {UpSetR} package^44^. The list of substrates for each SLC transporter was obtained from the SLC tables at slc.bioparadigms.org^25^. The list of tight junction genes was downloaded from the AMIGO gene ontology website^45,46^, using the Gene Ontology term GO:0070160.

### MeDP-seq bioinformatic processing

As MeDP-seq data is conceptually similar to ChIP-seq data, MeDP-seq reads were processed using the nf-core/chipseq pipeline^37^ version 1.2.1 on Ensembl reference genome Sscrofa11.1^39^. CpG methylation ratios were deduced from MeDP-seq coverages using BayMeth^47^ and data from the artificially methylated control samples. Genomic regions with anomalous coverage in input samples were identified using the {greylistchip} bioconductor package^48^ version 1.28.1 applied on each of the four input samples separately, and the four greylist regions were merged together. Genomic regions within our greylist were removed from the rest of the analysis.

To define lowly methylated regions (LMRs) we used the following steps: average DNA methylation ratio was computed for each sample in 100 bp windows. We then kept the 100 bp windows for which the DNA methylation ratio was under 40% in at least two samples within LRFI fed, LRFI fasted, HRFI fed, and HRFI fasted pigs. Consecutive 100bp windows were merged, and only genomic regions of 500 bp or more were kept. Unmethylated regions separated by 300 bp or less were merged. This resulted in the definition of 60,509 LMRs covering 0.03% of the genome (mean size: 1089 bp, median: 700 pb, minimum size: 500 bp, maximum size: 33,500 bp).

MeDP-seq coverage at LMRs was obtained for each sample using {EnrichedHeatmap} `normalizeToMatrix` function^49^. We used {DESeq2}^50^ to identify LMRs with differential coverage between the two LRFI input control samples and the two HRFI input control samples. This resulted in the identification of 9,572 LMRs with differential coverage between inputs from the two lines at an unadjusted p-value threshold of 5%. These regions were filtered as they might reflect structural genomic differences between the two lines resulting in misestimations of their DNA methylation state by MeDP-seq. Differentially methylated regions (DMRs) between the two lines were identified within the resulting 50,919 lowly methylated regions using {DESeq2}^50^ with a linear model taking into account the pig line (LRFI or HRFI) and condition (fed or fasted). Fed/fasted DMRs were called in each pig line independently using the same approach. In addition to an FDR adjusted p-value below 0.05 and after manual visualisations of some regions, we selected only regions with an absolute log2 fold change greater than 1 as differentially methylated regions in an attempt to further reduce false positives. DNA methylation ratio at differentially methylated regions were visualised using {epistack}^51^.

## Data and script availability

RNA-seq and MeDP-seq reads are available through the ENA database, id PRJEB46060 / ERP130249. Scripts used to process and analyse RNA-seq and MeDP-seq reads are available through a public gitlab repository at the address: forgemia.inra.fr/genepi/analyses/rosepigs. Gene expression counts, lists of differentially expressed genes, sample metadata tables, and DMR positions are available in this repository (forgemia.inra.fr/genepi/analyses/rosepigs/-/tree/master/processed_files).

## Results

### The duodenum transcriptome is different between the high- and low-feed efficiency pig lines

To compare pigs from the 10th generation of selection, 12 animals for each line were taken from 3 litters in the same breeding batch trying to balance sex ratios (figure 1A). After overnight fasting with unrestricted water access, feed was reintroduced to 6 HRFI and 6 LRFI animals in a single pen (balancing litters and sex, figure 1B). Duodenal mucosa samples were then collected, between 2 to 3 hours after feed re-introduction, or between 12 to 14 hours of feed restriction. Duodenal transcriptomes were obtained by RNA-sequencing after poly-A purification, enriching for messenger RNA. We detected 1,106 genes differentially expressed between the LRFI and HRFI lines, independent of the feeding status of the animals (figure 2A, supplementary table 4). In detail, 464 genes were identified as upregulated in the LRFI line, and 642 genes were identified as up-regulated in the HRFI line.

**Figure 2:**
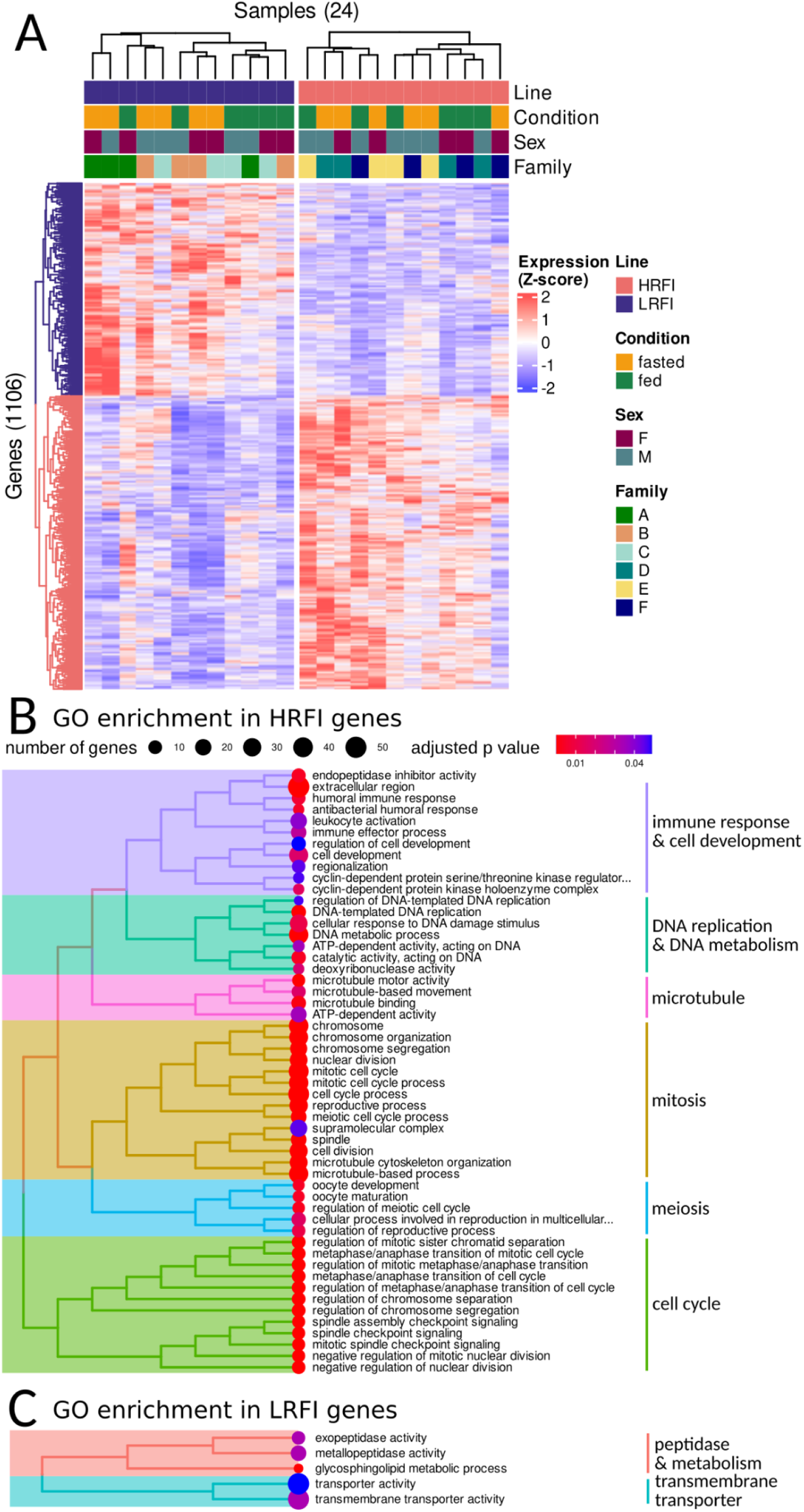
Differential gene expression between the duodenum of LRFI and HRFI lines. **A**. 1,106 genes were detected differentially expressed between LRFI and HRFI, including 464 genes upregulated in LRFI (top) and 642 genes upregulated on the HRFI line (bottom). **B & C**. Gene Ontology - Biological Process (GO-BP) enrichment analysis. GO-BP terms statistically enriched in the 642 genes upregulated on the HRFI line (**B**) and the 464 genes upregulated in LRFI line (**C**) are displayed as a tree using Jaccard similarity between each pair of GO-BP.

Functional enrichment analysis revealed that genes upregulated in HRFI were notably involved in various aspects of cell division, including spindle checkpoint signalling, chromosome segregation, DNA-templated DNA replication, and immune response (figure 2B). Fewer processes were functionally enriched in genes upregulated in the LRFI lines (figure 2C), including glycosphingolipid metabolic process and transmembrane transporter activity.

### The pig duodenum transcriptome response to feed intake is stronger in LRFI pigs than in HRFI pigs

For each line, we compared duodenal transcriptomes in fasted or fed pigs. We detected 2,222 differentially expressed genes in the LRFI pig line, but only 61 differentially expressed genes in the HRFI line (figure 3, A&B, supplementary table 4), leading to a total of 2,225 genes affected by feed intake in one or both pig lines. Visualisation of expression profiles revealed that in the LRFI line, all 6 fasted pigs were showing stark differences of expression as compared to the 6 fed pigs in the differentially expressed genes, while 3 fasted HRFI pigs and 2 fed HRFI pigs did not have a transcriptomic signature matching the feeding status signature of the LRFI line (figure 3A).

**Figure 3:**
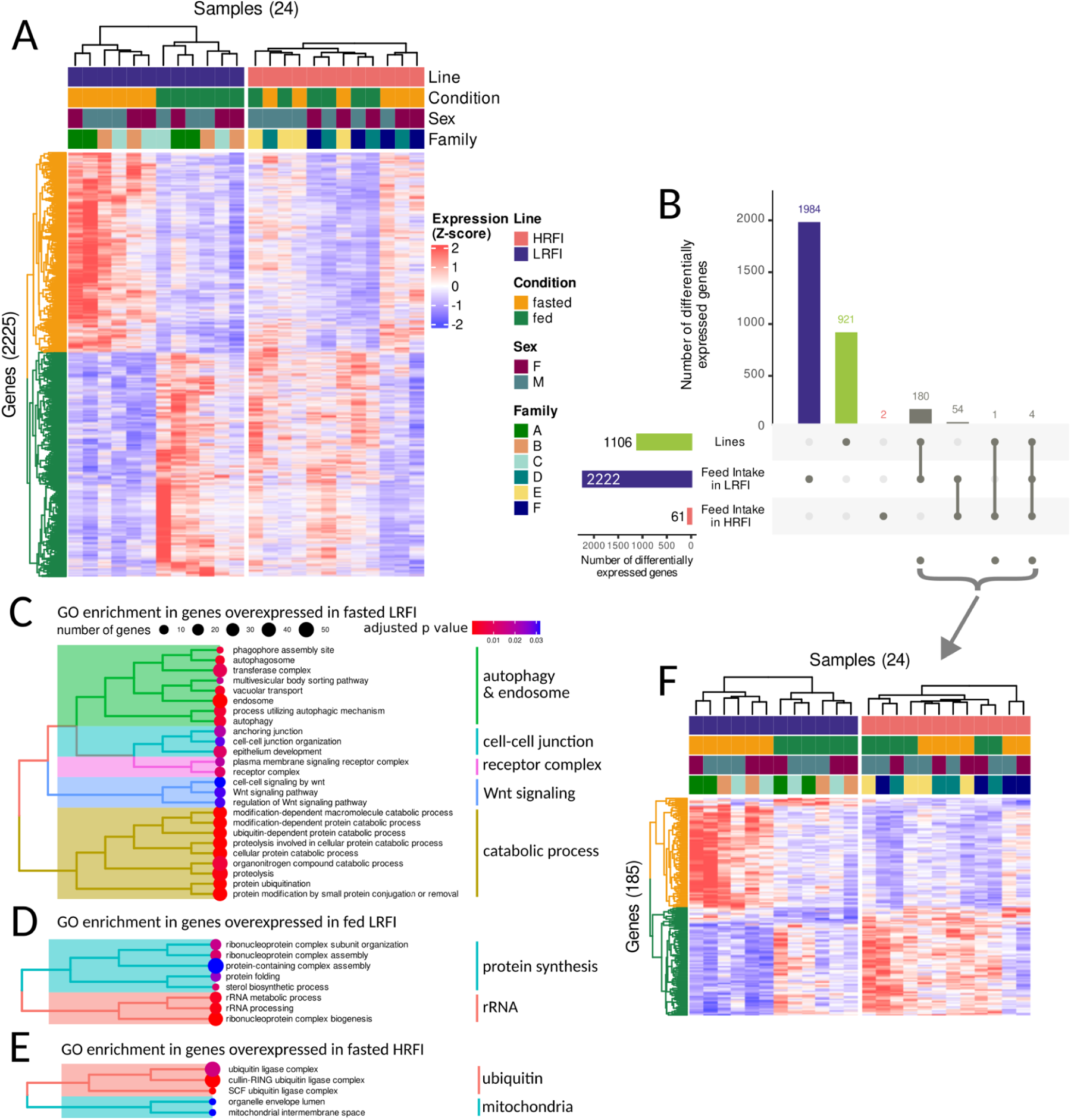
Duodenum transcriptomic response to feed intake. **A**. Gene expression heatmap of the 2,225 genes differentially expressed by feed intake in LRFI (2,222 genes), in HRFI (61 genes, including 58 genes also affected by feed intake in LRFI). **B**. Upset diagram showing the overlap between differentially expressed gene lists, between LRFI and HRFI (green), and due to feed intake in LRFI pigs (blue) or HRFI pigs (red). **C, D & E**. Gene Ontology - Biological Process (GO-BP) enrichment analysis. GO-BP terms statistically enriched in the 1,050 genes upregulated upon fasting in the LRFI line (**C**), the 1,172 genes upregulated in fed pigs from the LRFI line (**D**), and the 41 genes upregulated in fed pigs from the HRFI line (**E**) are displayed as a tree using Jaccard similarity between each pair of GO-BP. **F**. Gene expression heatmap of the 185 genes differentially expressed both by line and by feed intake in LRFI and/or HRFI pigs.

In the LRFI line, the 1,050 genes overexpressed in the fasted state were enriched in GO-BP categories linked to autophagy, cell junction and plasma membrane receptors, Wnt signalling pathway, and protein catabolic processes (figure 3C). The 1,172 genes overexpressed in the fed state in LRFI pigs were enriched in GO-BP categories linked to protein folding, ribosome biogenesis and the sterol biosynthetic process (figure 3D). In the HRFI line, the 41 genes overexpressed in the fasted state were enriched in GO-BP categories linked to ubiquitin ligase complex, and with mitochondrial envelope to a lesser extent (figure 3E). No GO-BP category was detected as enriched in the 20 genes overexpressed in the fed states in HRFI pigs.

Among the genes impacted by the feeding status in one or both lines, 185 genes were also detected as differentially expressed between the two lines (figure 3B). Visual inspection of these genes revealed that they mostly respond to feed intake in the LRFI line and not in the HRFI line, resulting in differences in average expression levels between the lines (figure 3F).

### Duodenal expression of nutrient transporter and tight junction genes was altered by the divergent selection on feed efficiency

The transmembrane transporter activity ontology term is overrepresented in genes with higher expression in LRFI than in HRFI (figure 2C). The Solute Carrier (SLC) gene family encodes for transmembrane transporters, including nutrient transporters expressed in the intestinal epithelium^25,26^. We therefore focused our analysis on the expression patterns of the 364 annotated SLC genes in the pig genome in our experimental setup (figure 4A). In the duodenum mucosa, 28 SLC genes are differentially expressed between the LRFI and HRFI lines: 17 are more expressed in the LRFI line, such as the folate transporter *SLC25A32*, 11 are more expressed in the HRFI line such as the glucose and galactose transporter *SLC2A10*. Thirty-five SLC genes are differentially expressed in the duodenum between fasted and fed LRFI pigs between: 18 are more expressed in fasted pigs, such as monosaccharides transporters *SLC5A1* and *SLC2A2* and the aspartate and glutamate transporter *SLC25A13*, 17 are more expressed in fed pigs, such as the glutamate transporter *SLC17A8*. No SLC gene was detected as differentially expressed by feed intake in the duodenum of HRFI pig.

**Figure 4:**
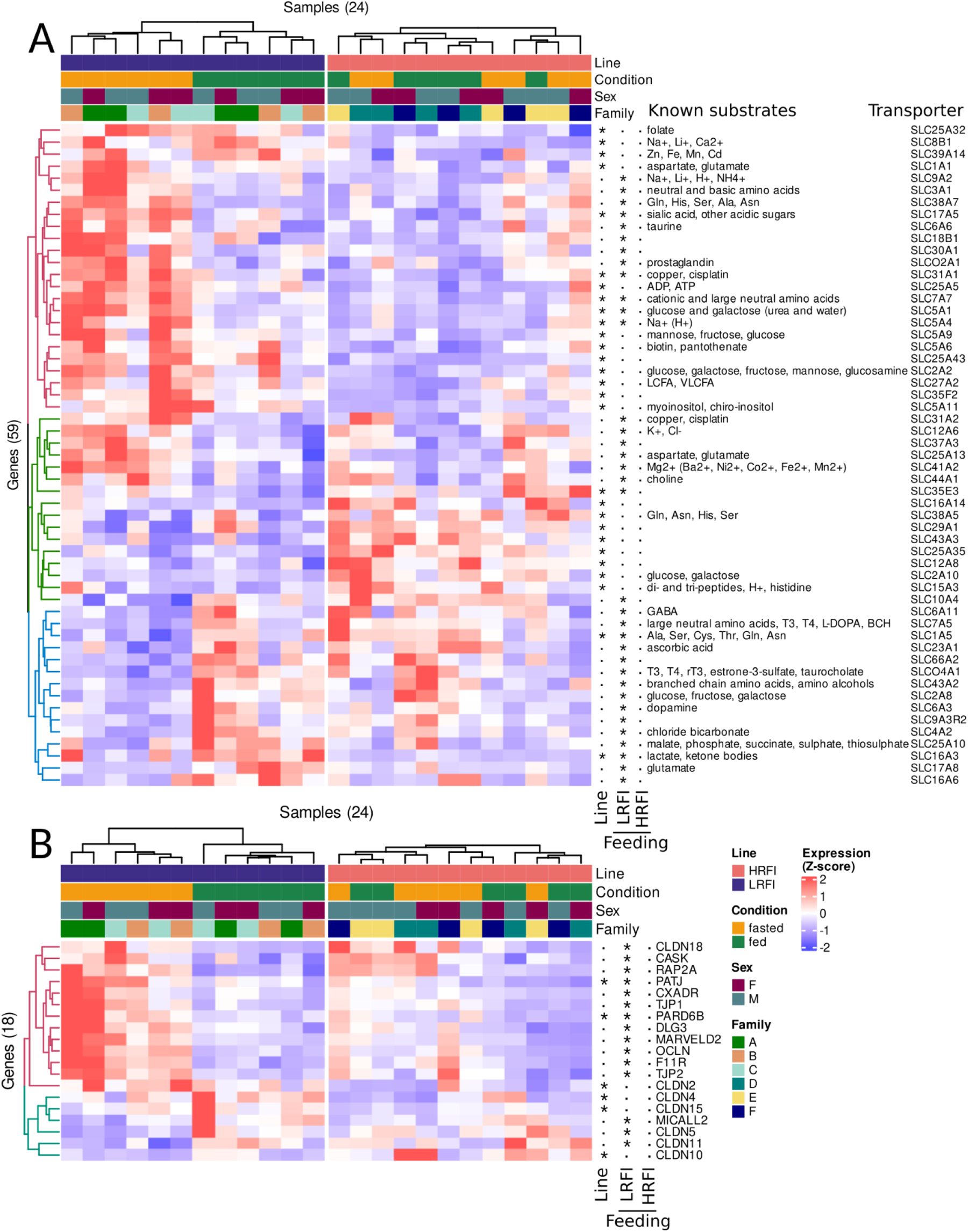
Gene expression heatmaps of the 59 *SLC* genes (A) or 18 tight junction genes (B) that are differentially expressed either between the LRFI and HRFI lines, or by feed intake in each line. Transporter names and known substrates are noted on the right of the heatmap. Three symbolic columns indicate if each gene is significantly differentially expressed (marked with “*”) or not (marked with “”) between the two lines (Line column), or by feed intake in the LRFI line (LRFI column), or by feed intake in the HRFI line (HRFI column). Side trees are coloured according to hierarchical clustering, and are only used as visualisation guides.

The cell-cell junction organisation ontology term was enriched in genes with higher expression in fasted LRFI pigs than in fed LRFI pigs (figure 3C). We therefore focused our analysis on the expression patterns of the 77 annotated tight junction genes in our experimental setup (figure 4B). In the duodenum mucosa, 5 tight junction genes are differentially expressed between the LRFI and HRFI lines, and all are more expressed in the LRFI line: *Claudin-2* (*CLDN2), CLDN4, CLDN15, Partitioning defective 6 homolog beta* (*PARD6B*) and *Protein Associated to Tight Junctions* / *InaD-like protein* (*PATJ*). Fifteen tight junction genes are differentially expressed by short term feed intake in the duodenum of LRFI pigs: 12 are more expressed during short term fasting, such as *Occludin* (*OCLN*), *Junctional adhesion molecule A* (*F11R*) or *PATJ*, and 3 are more expressed after feed intake: *CLDN5, CLDN11* and *MICAL-Like 2* (*MICALL2*). No tight junction gene was detected as differentially expressed by feed intake in the duodenum of HRFI pig.

### Differences in DNA methylation profiles in the duodenum of LRFI and HRFI pigs

DNA methylation profiles were obtained by MeDP-seq on the same duodenum samples. Although MeDP-seq primarily measures DNA methylation density, we converted it into DNA methylation ratio using fully methylated control samples (see method)^47^. For all samples, DNA methylation was lower near the transcription start site (TSS) of expressed genes than at the TSS of un-expressed genes (supplementary figure 1). We detected low DNA methylation regions (LMRs) in each experimental group as putative regulatory regions. Amongst the 50,872 LMRs, 409 were differentially methylated between LRFI and HRFI, 26 were detected as differentially methylated between fasted and fed LRFI pigs, and 3 were detected as differentially methylated between fasted and fed HRFI pigs (figure 5, supplementary table 5, 6, and 7). We did not find global associations between DMRs and differential gene expression, but some overlaps were intriguing. Of the 409 HRFI and LRFI DMRs, 20 were within 20kb of a differentially expressed gene between LRFI and HRFI pigs (supplementary table 8), and the associations between changes in DNA methylation and changes in gene expression were not always in the same direction. Amongst those were *CLDN10* (hypomethylated on the first intron in HRFI, and more expressed in HRFI than in LRFI, figure 6 A&B), *PHGDH* (Phosphoglycerate dehydrogenase, hypomethylated in an internal exon and intron in HRFI, and more expressed in HRFI than in LRFI, figure 6 C&D), *ITLN2* (Intelectin-2, hypomethylated in an internal intron in LRFI, and more expressed in HRFI than in LRFI, figure 6 E&F), *FER* (Proto-oncogene tyrosine-protein kinase, hypomethylated in an internal intron in HRFI, and more expressed in LRFI than in HRFI, figure 6 G&H), and the long non-coding RNA *ENSSSCG00000044004* (hypomethylated at its promoter in HRFI, and more expressed in HRFI than in LRFI, figure 6 I&J). Of the 29 DMRs between fed and fasted pig in one of the lines, 3 were within 20kb of a differentially expressed gene between fasted and fed LRFI pigs (supplementary table 9). Amongst those were *KDSR* (3-dehydrosphinganine reductase, hypomethylated in fed LRFI pigs compared to fasted LRFI pigs, and less expressed in fed LRFI pigs compared to fasted LRFI pigs, figure 6 K&L). KDSR3 was also less expressed in fed HRFI pigs than in fasted HRFI pigs, but to a lower extent than in LRFI pigs.

**Figure 5:**
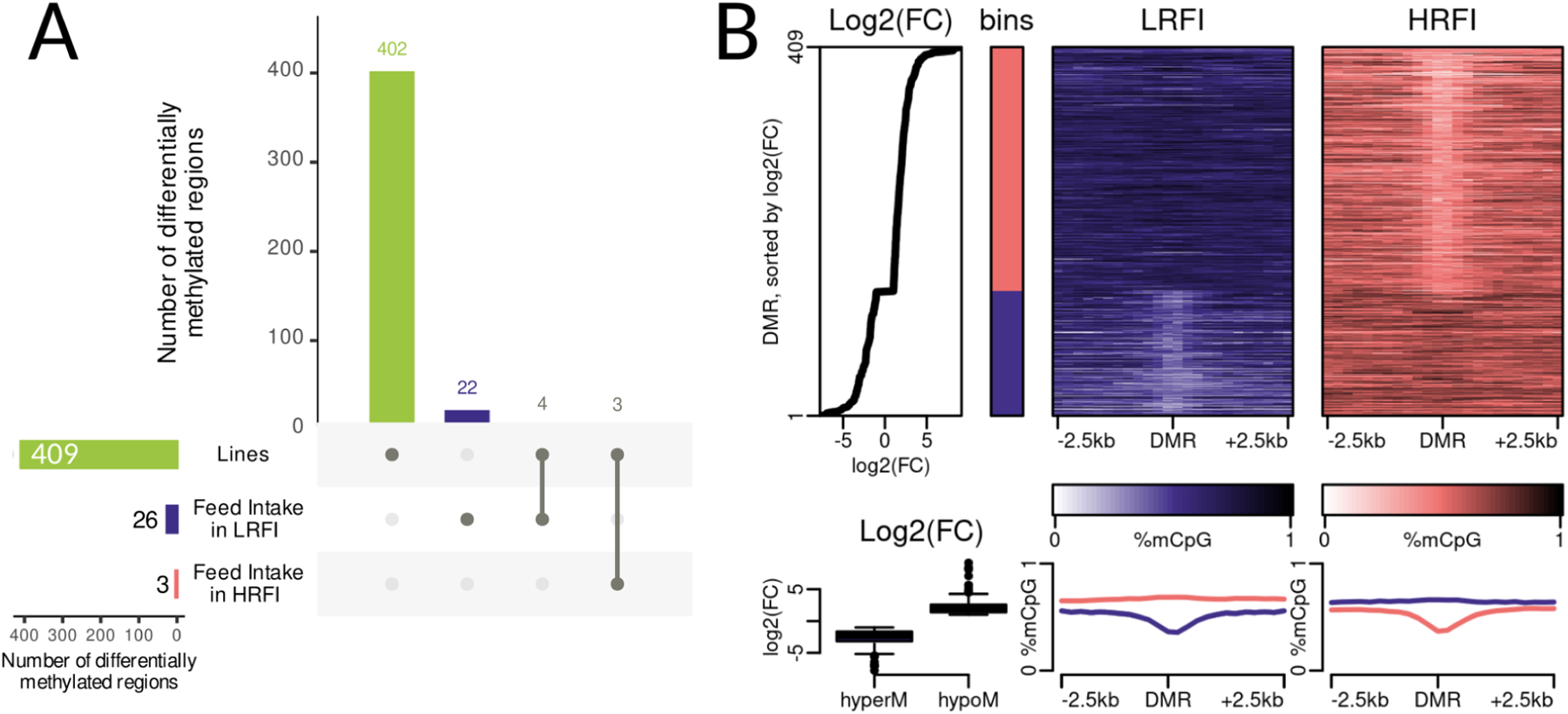
Differentially methylated regions in the duodenum between LRFI and HRFI samples. **A**. Upset plot of differentially methylated regions. Green: between LRFI and HRFI samples. Blue: between fasted and fed samples in LRFI pigs. Red: between fasted and fed samples in HRFI pigs **B**. DNA methylation stack profiles at differentially methylated regions between LRFI and HRFI samples. Top: log2 Fold Change (FC) of MeDP-seq normalised coverage at DMR. The 409 LRFI/HRFI DMRs are sorted according to their fold change. Then: each DMR is categorised into a hypermethylated in LRFI bin (hyperM, red) or hypomethyated in LRFI bin (hypoM, blue). Then: Average DNA methylation ratio at ± 2.5kb of DMR centers, in LRFI (blue) or HRFI (red) samples. Bottom: Distribution of fold changes in each bin, then average DNA methylation ratios in each bin (hyperM in LRFI: red, hypoM in LRFI: blue).

**Figure 6:**
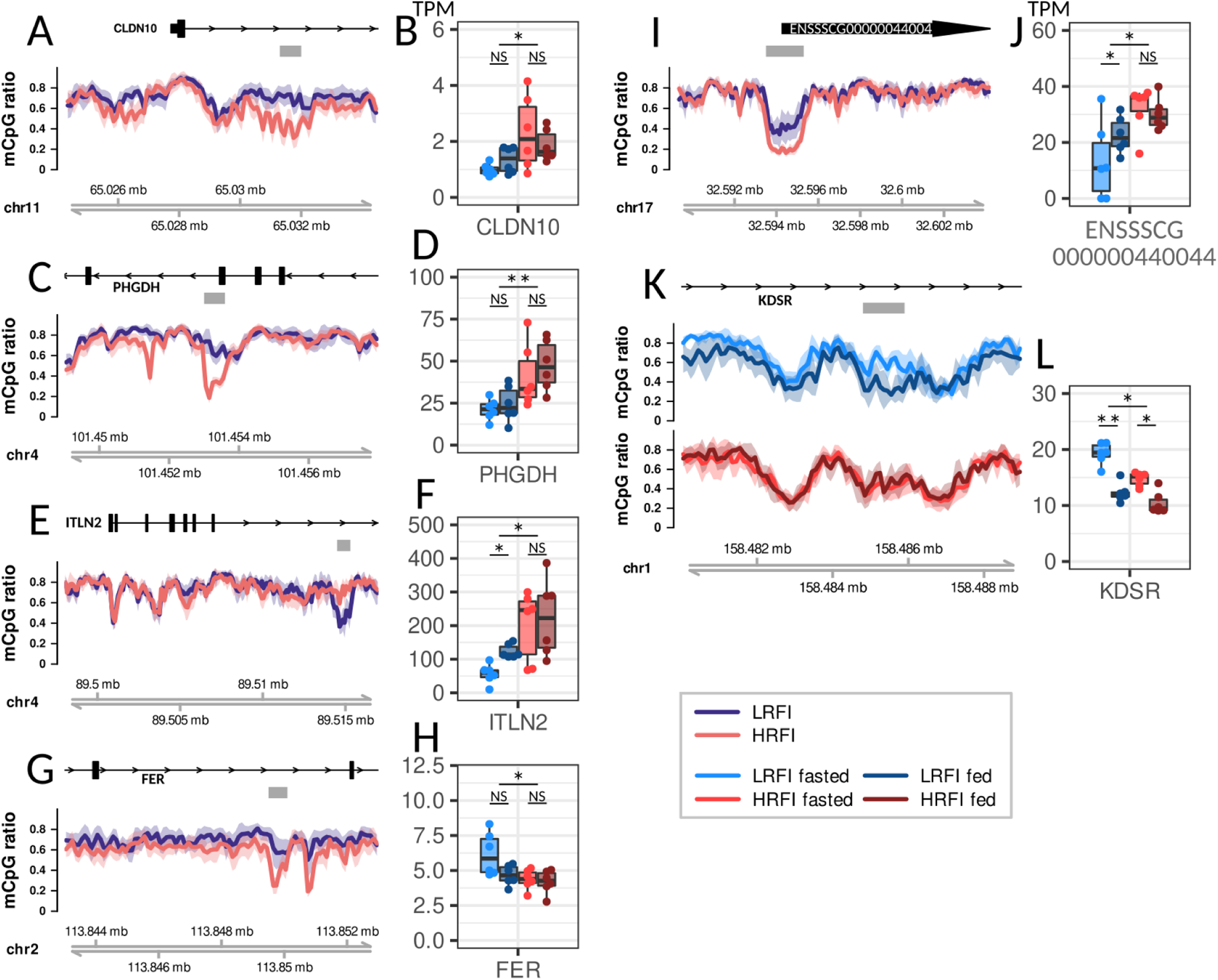
Some differentially methylated regions at proximity differentially expressed genes. **A**,**C, E, G, I, K**: DNA methylation profiles around differentially methylated regions. From top to bottom: gene structure in black. Thick boxes are exons, thin lines are introns, arrows indicate gene direction. Grey box indicates DMR location. Then, average DNA methylation ratio profiles ± 95% confidence interval per line (**A, C, E, G, I**, blue: LRFI, red: HRFI) or per condition in each line (**K**, light blue: LRFI fasted, dark blue: LRFI fed, light red: HRFI fasted, dark red: HRFI fed). Then genome coordinates of the corresponding region. **B, D, F, H, J, L**: Gene expression values in LRFI fasted (light blue), LRFI fed (dark blue), HRFI fasted (light red), HRFI fed (dark red) samples. Unit in transcript per million (TPM). Each dot is a sample, and boxplots are added to ease visualisation. NS: non significant. *:FDR-adjusted p-values < 0.05. **: FDR-adjusted p-values < 0.01. Adjusted p-values are from the differential expression analysis using *limma voom*.

## Discussion

Here we demonstrated that the divergent selection on feed efficiency changed the duodenum mucosal transcriptome in pigs, identifying 1,106 genes differentially expressed between the LRFI and HRFI lines. Genes overexpressed in LRFI were enriched in gene ontologies relevant to glycosphingolipid metabolic process, transmembrane transport, and exopeptidase and metallopeptidase activity. These functions do not directly mirror results from the transcriptome of muscle, liver and adipose tissues^12^, or in blood, muscle and liver metabolism^13^. Therefore, it is likely that genetic differences due to selection have led to differences in gene expression that are distinct between tissues, and are not systematic. Functions enriched in HRFI pigs were overwhelmingly related to mitosis related processes, such as chromosome separation, DNA replication, and mitosis checkpoint. It is not known if it is due to issues with cell division in the HRFI line that could be frequently failing, or if it simply reflects a higher cell division rate in the duodenum mucosa of HRFI pigs. Histological examination of the duodenum from HRFI and LRFI pigs might be very informative, especially if coupled with measures of cell divisions.

The LRFI line shows a very strong transcriptomic response to feed intake in its duodenum mucosa, with 2,222 differentially expressed genes. Gene expressed after fasting were enriched in catabolic functions and autophagy, while genes expressed after feed intake were more anabolic, which was perhaps to be expected. The large number of differentially expressed genes between fasted and fed LRFI pigs is in contrast to the weak response observed in HRFI pigs, with only 85 differentially expressed genes. It is not known if selection has increased the transcriptomic response in the LRFI line, suppressed the transcriptomic response in the HRFI line, or both at the same time. In mice, only a modest transcriptomic response to feed intake was observed by Yoshioka et al.^22^, but it might be due to a relatively poor sensitivity of the SAGE method used in the mice study. We observed that several nutrient transporters had their expression level increased or decreased by feed intake in the LRFI line. We notably confirmed in pigs the previous observation of the downregulation of *Slc5a1* in fasted mice^22^. *SLC5A1* encodes for the glucose transporter SGLT1. Different glucose absorption dynamics between LRFI and HRFI lines might lead to different insulinemia, as it has been observed in the same pig lines in earlier generations^13,14^. *SLC4A2* was more expressed in fed than in fasted LRFI pigs. It encodes an anion transporter involved in balancing the cellular pH^52^. It was also recently identified in a Epigenome Wide Association Study (EWAS): DNA methylation profiles located at *SLC4A2* correlated with food fussiness in children^53^. We did not find any DMR near *SLC4A2* in our dataset. More generally, it is tempting to hypothesise that the lack of transcriptomic response in the duodenum between fasted and fed HRFI pigs might explain some feeding behaviours observed in the HRFI line, for example due to their putative reduced nutrient sensing.

Genes from the cell-cell junction organisation ontology were over-represented in genes more expressed in the fasted condition in the LRFI line. Cell-cell junction, tight junctions in particular, allow the intestine epithelium to act as a barrier between the intestinal lumen and the rest of the organism^54^. It may be that LRFI pigs increase the sealing of tight junctions between meals, but HRFI pigs do not. Additional analyses will be required to test this hypothesis, notably histological analyses of the duodenum mucosa or by measuring duodenal permeability. The pathophysiological consequences of this regulation of tight junction induced by genetic selection for feed efficiency should also be investigated, notably in relation with the susceptibility to enteric infections^55,56^.

DNA methylation tends to be absent from active genomic regions^57^. Using MeDP-seq, we detected 409 differentially methylated regions between LRFI and HRFI lines using conservative filtering steps. It is likely that a finer DNA methylation measure such as whole genome bisulfite conversion followed by sequencing would have allowed the identification of more differentially methylated regions. Nonetheless, it can be concluded that the selection process for feed efficiency has resulted in distinct duodenal DNA methylation profiles. We only detected 26 and 3 differentially methylated regions between fasted and fed pigs of the LRFI and HRFI line, respectively. Thus, while short term feed intake drastically changes the gene expression profiles of the duodenal mucosa, it barely affects the DNA methylation profiles. Therefore, most**—**if not all**—**duodenum gene expression changes between fasted and fed samples were not mediated by DNA methylation changes. It is likely that changes in DNA methylation profiles need more time than changes in gene expression profiles. Nonetheless, it appears that for DNA methylation profiles as for gene expressions, the LRFI line is more sensitive than the HRFI line to meal intake.

While we could not correlate all the changes in DNA methylation profiles with overall changes in gene expressions, we identify several examples of DMRs located near or within differentially expressed genes (figure 6). This is the case for example for *CLDN10*, involved in tight junctions, *PHDGH*, involved in serine biosynthesis, *ITLN2*, coding for a carbohydrate binding protein, *FER*, a tyrosine kinase regulating cell junctions, *MUC13* (data not shown), a mucin gene, or *KDSR*, involved in the sphingolipid metabolism. The associations between DNA methylation changes and gene expression changes could be either positive or negative, which has been previously reported^58^. With this dataset alone, we cannot decipher causalities between the changes in DNA methylations and changes in gene expression levels. Differences between the lines are likely due to genetic effects, but we cannot discriminate if the same genetic differences are involved in changes in DNA methylation and gene expressions, or if the differences we observed are simply coincidental. The long non-coding RNA gene *ENSSSG00000044004* has a DMR right at a lowly methylated region on its promoter, and more DNA methylation is associated with a reduction of its expression. For this case, the correlation between DNA methylation changes and gene expression changes likely reflects causality links. This uncharacterised long non-coding RNA lies between the vasopressin and the oxytocin genes, for which the synteny is conserved in most vertebrates^59^.

The pig is thought to be a good model of human digestive physiology^60–62^, due to its closest diet, diurnal rhythm and size when compared to rodent models. The dataset produced here might prove useful to better understand the transcriptomic response to feed intake of the human duodenum. Therefore, our dataset is publicly available, at the sequencing read levels, in the form of gene expression tables, and in lists of differentially expressed genes. In addition, better understanding of the biological mechanisms of feed efficiency may contribute to improvements of feed efficiency in pig farming, leading to a more sustainable production.

## Acknowledgment

The authors would like to thank Sophie Leroux and all the GenESI staff for their help with the pig experimentation and sampling^35^. We are grateful to the GenoToul bioinformatics platform Toulouse Occitanie (Bioinfo GenoToul^63^) for providing computing and storage resources. GD thanks Juliette Riquet for her curiosity about tight junction proteins, Sylvain Foissac for his insight in lncRNA gene annotations, and the Bioinfo-fr IRC/Discord community for their support and insight on Gene Ontology analysis.

## Acronyms

DMR: Differentially methylated regions
GO-BP: Gene Ontology - Biological Process
HRFI: Pig line selected for a High RFI (low feed efficiency)
LMR: Lowly methylated regions
LRFI: Pig line selected for a low
RFI: (high efficiency)
MeDP-seq: DNA methylation precipitation followed by high throughput sequencing
RFI: Residual Feed Intake
SAGE: Serial analysis of gene expression
SLC: Solute Carrier, a gene family of transmembrane nutrient transporter
TPM: Transcript per million, a normalised expression value.
TSS: Transcription Start Site

## Supplementary figure

**Supplementary figure 1:**
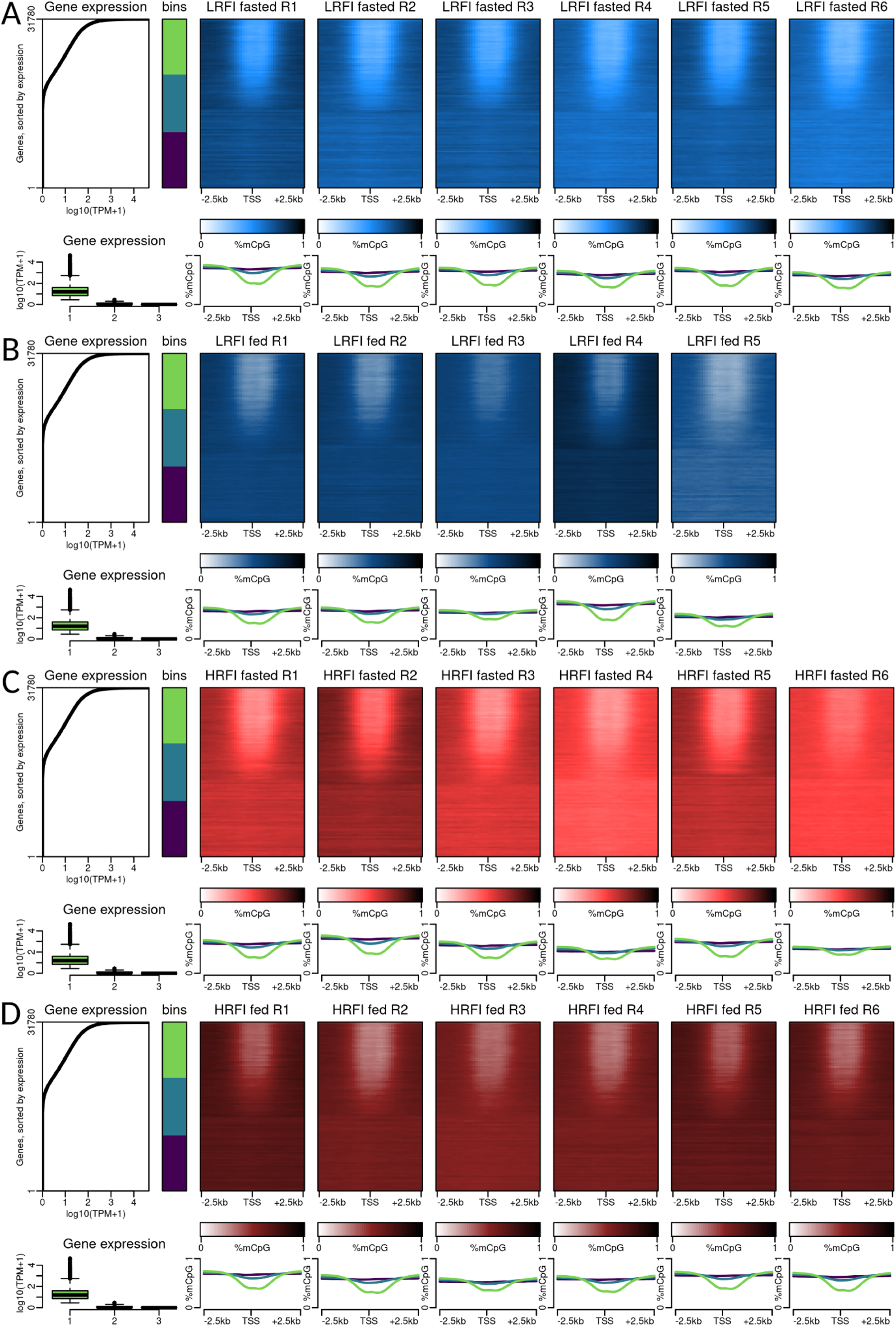
Low DNA methylation at the promoter of expressed genes. In each MeDP-seq sample, we sorted genes according to expression levels from their corresponding RNA-seq sample, from top (high expression) to bottom (low expression). Genes were binned in three groups of equal size : light green (high expression), dark green (low to no expression), and purple (no expression). For each gene, DNA methylation ratio is represented in the -2.5kb +2.5kb window surrounding their Transcription Start Site (TSS). Bottom panels display the expression values in each bin of genes, as well as their average DNA methylation ratio around TSS. **A**. LRFI samples in the fasted condition. **B**. LRFI samples in the fed condition (one sample failed during library preparation). **C**. HRFI samples in the fasted condition. **D**. HRFI samples in the fed condition.

